# Early Dietary Exposures Epigenetically Program Mammary Cancer Susceptibility through IGF1-mediated Expansion of Mammary Stem Cells

**DOI:** 10.1101/2020.11.15.383570

**Authors:** Yuanning Zheng, Linjie Luo, Isabel U. Lambertz, Robin Fuchs-Young

**Affiliations:** Department of Molecular and Cellular Medicine, Texas A&M Health Science Center, Bryan, TX 77807, USA

**Keywords:** Diet, Insulin-Like Growth Factor I, Epigenetics, Stem Cells, Carcinogenesis

## Abstract

Dietary exposures at early developmental stages have been shown to program lifetime breast cancer susceptibility. We previously reported that manipulation of gestational and postweaning diets leads to different mammary tumor outcomes in carcinogen-treated mice. The high tumor incidence (HT) groups (average 61.5% tumor incidence) received a low-fat, low-sugar, mildly restricted (12%v/v) (DR) diet during gestation, followed by a high-fat, high-sugar (HF) diet postweaning. Conversely, the low tumor incidence (LT) groups (average 20% tumor incidence) received the HF diet during gestation, followed by the DR diet postweaning. Herein, we extended these findings by demonstrating that HT animals had an expanded mammary stem cell (MaSC) population compared to LT animals before puberty, and this expansion persisted into adulthood. IGF1 expression was increased in mammary stromal cells from HT animals, which promoted the self-renewal capacity of MaSCs in a paracrine fashion. This increased IGF1 expression was programmed prepubertally through DNA hypomethylation of the IGF1 promoter 1, mediated by decreased DNMT3b levels. IGFBP5 mRNA and protein levels were also reduced in mammary tissues from HT animals, indicating an increased bioavailability of tissue IGF1. In association with these changes, mammary tissues from carcinogen-treated HT animals developed an increased proportion of mammary adenosquamous carcinomas compared to carcinogen-treated LT animals. This study provides novel mechanistic insights into how early dietary exposures program mammary cancer risk and tumor phenotypes by increasing IGF1 expression through epigenetic alterations, thereby expanding the MaSC population, resulting in a higher number of carcinogen targets susceptible to transformation in adulthood.

**Significance:** Early high-fat dietary exposure programs lifetime mammary cancer susceptibility before puberty through epigenetic alterations of IGF1 promoters and IGF1-mediated paracrine regulation of mammary stem cell homeostasis.

## Introduction

Environmental exposures, such as diet, are critical determinants of breast cancer risk, and recent studies indicate that exposures during the early developmental stages can program lifetime cancer susceptibility [1–3]. Data from Dutch Famine studies show that the breast cancer incidence is increased in women who were exposed to severe dietary restriction *in utero* or in childhood (age 2~9) during the 1944-1945 hunger winter in World War II compared to their unexposed counterparts [2,4]. Besides dietary restriction, early life high-fat diet exposure has also been shown to program mammary cancer susceptibility [5–7]. We previously reported that gestational exposure to a high-fat and high-sugar diet has a protective effect on mammary carcinogenesis in mice, whereas exposure to the high-fat and high-sugar diet postweaning increases the mammary cancer hazard ratio by 2 to 5.5 times [3]. Despite substantial efforts, the underlying mechanisms of how such early dietary exposures program mammary cancer risk remain unresolved.

Previous studies have linked dietary exposures to dysregulation of tissue stem and progenitor cell homeostasis [8,9]. Yilmaz *et al.* reported that mice fed a high-fat diet had an increased number of intestinal stem and progenitor cells compared to mice fed a low-fat control diet, and *in vivo* transplantation assays show that feeding mice a high-fat diet primes intestinal progenitors for transformation via activation of the PPAR δ signaling pathway [8]. The mammary gland contains multipotent mammary stem cells (MaSCs) and lineage-committed progenitors [10,11], and both mammary stem and progenitor cells can be cancer-initiating cells [12–14], with the lifetime risk for developing mammary cancer positively associated with the number of these cells in the mammary epithelium [3,14–16]. *In vivo* labeling studies show that mammary stem and progenitor cells have increased longevity compared to differentiated cells, suggesting that they can be primary susceptible targets that accumulate genetic mutations leading to mammary carcinogenesis [17,18]. Morel *et al.* reported that Ras-transformed human MaSCs have a reduced accumulation of reactive oxygen species during cell proliferation compared to Ras-transformed differentiated cells [12]. Therefore, MaSCs have increased survival advantages after oncogenic insults, making them potential tumor-initiating cells [12].

Epigenetic alteration is a critical molecular mechanism underlying the effects of early environmental exposures on programming cell differentiation, proliferation and metabolism (reviewed in [19,20]). The major components of epigenetic processes include DNA methylation, histone modifications and small non-coding RNAs. Disruption of these epigenetic processes is implicated in cancer initiation and progression in many organs, including the breast (reviewed in [21]). Therefore, we hypothesized that early dietary manipulations alter the number of MaSCs at an early stage of development through epigenetic alterations, thereby affecting the lifetime mammary cancer risk.

To investigate this hypothesis, we used the cross-over feeding mouse model established in our previous study [3]. Outbred SENCAR mice were fed for ten weeks with either a high fat, high sugar (HF) diet *ad libitum* to induce metabolic syndrome, or a defined low fat, low sugar and mildly restricted (12%v/v) diet (DR) used as control. These mice are then bred, and cross-over feeding continues during three critical time windows of development: gestation, lactation and postweaning. We have shown that manipulations of gestation and postweaning diet significantly affect mammary tumor susceptibility [3]. As shown in **Table 1 (** adapted from ref. [3]), high-tumor susceptible (HT) animals (groups HT-A and HT-B) were exposed to a maternal DR diet during gestation, followed by either a DR or HF diet during lactation, and a HF diet postweaning (DR/ _/HF). Conversely, the low-tumor susceptible (LT) animals (groups LT-C and LT-D) were exposed to a HF diet during gestation, followed by either a DR or HF diet during lactation, and a DR diet postweaning (HF/_ /DR) [3]. In our previous work, we reported that animals from group HT-A have an increased proportion of mammary epithelial cells (MECs) with stem cell surface markers compared to group LT-C in 5-week old mammary tissues, indicating that these dietary manipulations may have affected MaSC homeostasis before puberty [3].

**Table 1.** The DR/_ /HF dietary regimens increase mammary tumor susceptibility

| Tumor Groups | Group | Gestation Diet | Lactation Diet | Post-Weaning Diet | Mammary Tumor Incidence |
| --- | --- | --- | --- | --- | --- |
| High Tumor (HT) | HT-A | DR | HF | HF | 65%<sup>a</sup> |
| High Tumor (HT) | HT-B | DR | DR | HF | 58%<sup>a</sup> |
| Low Tumor (LT) | LT-C | HF | HF | DR | 22%<sup>b</sup> |
| Low Tumor (LT) | LT-D | HF | DR | DR | 18%<sup>b</sup> |

(a ≠ b, Pearson's chi-squared test, $P < 0.03$ )

The purpose of the current study was to accurately enumerate and definitively characterize the MECs harboring stem cell markers, to delineate the mechanisms responsible for their expansion resulting from exposure to HT dietary regimens, and to define their role in increasing tumor risk. We show that the pro-tumorigenic DR/ _ /HF dietary regimens increased the expression of insulin-like growth factor 1 (IGF1) in mammary stromal cells from HT animals by inducing DNA hypomethylation of the IGF1 promoter 1. This increased stromal IGF1 expression promoted MaSC self-renewal in a paracrine fashion and expanded the MaSC population before the animals reached puberty. Our results demonstrate a novel mechanism through which dietary manipulations affect mammary tumor incidence by inducing specific epigenetic alterations at an early developmental stage, thereby changing the hierarchical organization of the glandular epithelium and increasing the number of susceptible carcinogen targets.

## Results

### The pro-tumorigenic dietary regimens of the HT groups expanded the MaSC-enriched compartment in both pre- and postpubertal female mice

To assess the effect of the dietary regimens on MaSC numbers, we analyzed MECs from 5-week old, prepubertal mice and 10-week old, postpubertal mice from both HT and LT groups (Table 1) by fluorescence-activated cell sorting (FACS). The MaSC-enriched population was identified by previously validated cell surface markers (Lin^−^/CD24^med^/CD29^hi^) with the gating strategy shown in Supplementary Fig. S1A [10,22]. In prepubertal mammary tissues, the proportion of MaSC-enriched cells from group HT-A was 3.5-times higher than in group LT-C and group LT-D (**Fig. 1A** and **B,** *P* < 0.0001). Similarly, the proportion of MaSC-enriched cells from group HT-B was 2-times higher compared to both LT groups (**Fig. 1A** and **B,** *P*<0.05). To determine whether this increase in the proportion of the MaSC-enriched population in HT animals persisted into adulthood, we also performed FACS analyses with postpubertal mammary glands. We found that the proportion of MaSC-enriched cells in glands from both HT groups was still increased by two times compared to the LT groups (**Fig. 1C** and **D,** *P*<0.01). These results indicate that exposure to the pro-tumorigenic DR/_ /HF dietary regimens expanded the MaSC-enriched compartment before puberty, and this expansion was maintained postpubertally.

**Figure 1.**
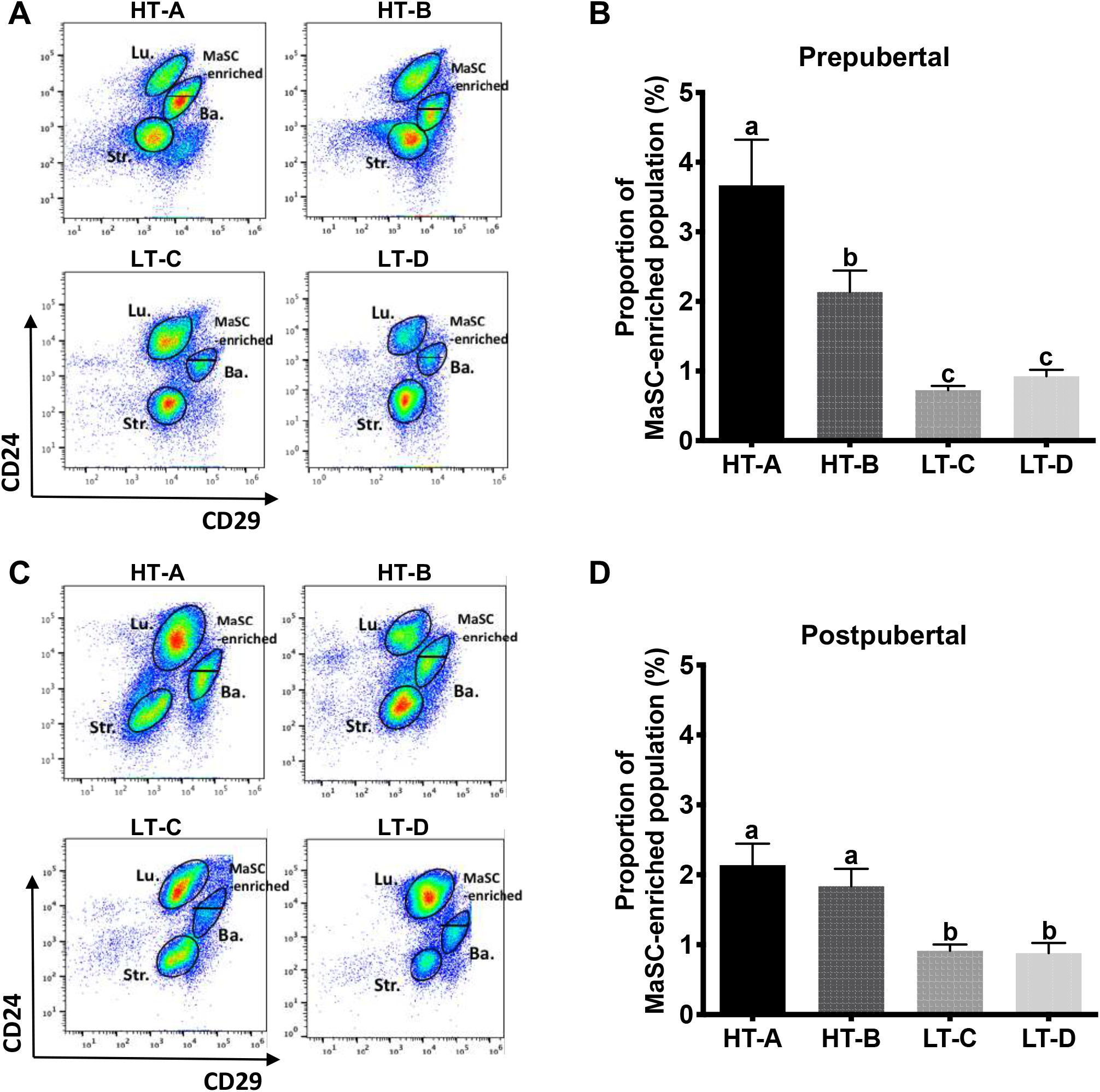
The pro-tumorigenic dietary regimens of the HT animals expanded the MaSC-enriched compartment in both pre- and postpubertal female mice. **(A)** Representative gating plots showing the MaSC-enriched population in 5-week old mammary glands. MaSC-enriched, basal (Ba.), luminal (Lu.) and stromal (Str.) cells were gated into Lin^−^CD24^med^CD29^hi^, Lin^−^ CD24^+^CD29^hi^, Lin^−^CD24^hi^CD29^low^ and Lin^−^CD24^low^CD29^low^ subpopulations, respectively. **(B)** Frequency of MaSC-enriched populations in mammary glands of 5-week old HT groups (HT-A & HT-B) and LT groups (LT-C & LT-D) animals (n≥8). **(C)** Representative gating plots showing the MaSC-enriched population in 10-week old mammary glands. **(D)** Frequency of MaSC-enriched populations in mammary glands of 10-week old HT groups and LT groups animals (n≥6). Mean ± SEM are shown. Differences were compared by one-way ANOVA. Pairwise comparisons were performed using Fisher’s LSD test. Groups are significantly different from each other if they do not share a letter (a≠b≠c, *P*<0.05).

Interestingly, when counting the total number of MECs dissociated from abdominal/inguinal mammary tissues, we found that prepubertal animals from group HT-A had twice the number of MECs compared to age-matched LT groups (Supplementary Fig. S1B). When multiplying the frequency of the MaSC-enriched population by the total number of MECs, the absolute number of cells with MaSC markers in prepubertal HT-A animals was 4.2-times higher than in age-matched LT groups (Supplementary Fig. S1C). In postpubertal glands, the number of MECs from both HT groups was 2-times higher compared to the LT animals, resulting in an average increase in the number of MaSC-enriched cells by 3.7 times in HT animals compared to the LT animals (Supplementary Fig. S1D and E). These results demonstrate that the absolute number of susceptible targets for mammary carcinogenesis was significantly higher in mammary tissues from HT animals than in LT animals.

### The number of mammosphere-initiating cells was higher in HT animals than in LT animals

Due to a lack of an exclusive marker for isolating MaSCs from Lin^−^/CD24^+^/CD29^hi^ basal cells, the FACS-sorted Lin^−^/CD24^med^/CD29^hi^ population cannot be used to directly enumerate MaSCs due to its compositional heterogeneity [10,22]. However, an intrinsic characteristic of MaSCs is their ability to self-renew and form sphere-like colonies (mammospheres) in non-adherent cultures [23]. The MaSC frequency in a given population of MECs is reflected by the number of mammospheres formed, particularly in the second and later serial passages [24,25]. In this study, we performed serial passages of mammospheres and compared the mammosphere-forming efficiency of MECs from HT and LT animals in the second (P2) passage. In addition, a mammosphere limiting dilution assay (LDA) was performed in the P3 passage to reliably quantitate MaSC frequency [25,26]. As shown in **Fig. 2A**, the number of P2 mammospheres formed using MECs from the prepubertal HT groups was significantly increased by 2.5 times in HT-A and 1.8 times in HT-B than those from the LT groups, respectively. P2 mammospheres were then disaggregated and plated in 96-well plates to perform an LDA of P3 mammospheres. In prepubertal mammary tissues, the frequency of MaSCs was 2.2-times higher in HT-A animals compared to LT-C (1 in 177 versus 1 in 404, *P*<0.001) and LT-D animals (1 in 177 versus 1 in 379, *P*<0.005) (**Fig. 2B** and Supplementary Table S1). Prepubertal HT-B animals also had a 1.5-time increase in MaSC frequency when compared to LT-C (1 in 254 versus 1 in 404, *P*<0.05) and LT-D animals (1 in 254 versus 1 in 379, *P*<0.05).

**Figure 2.**
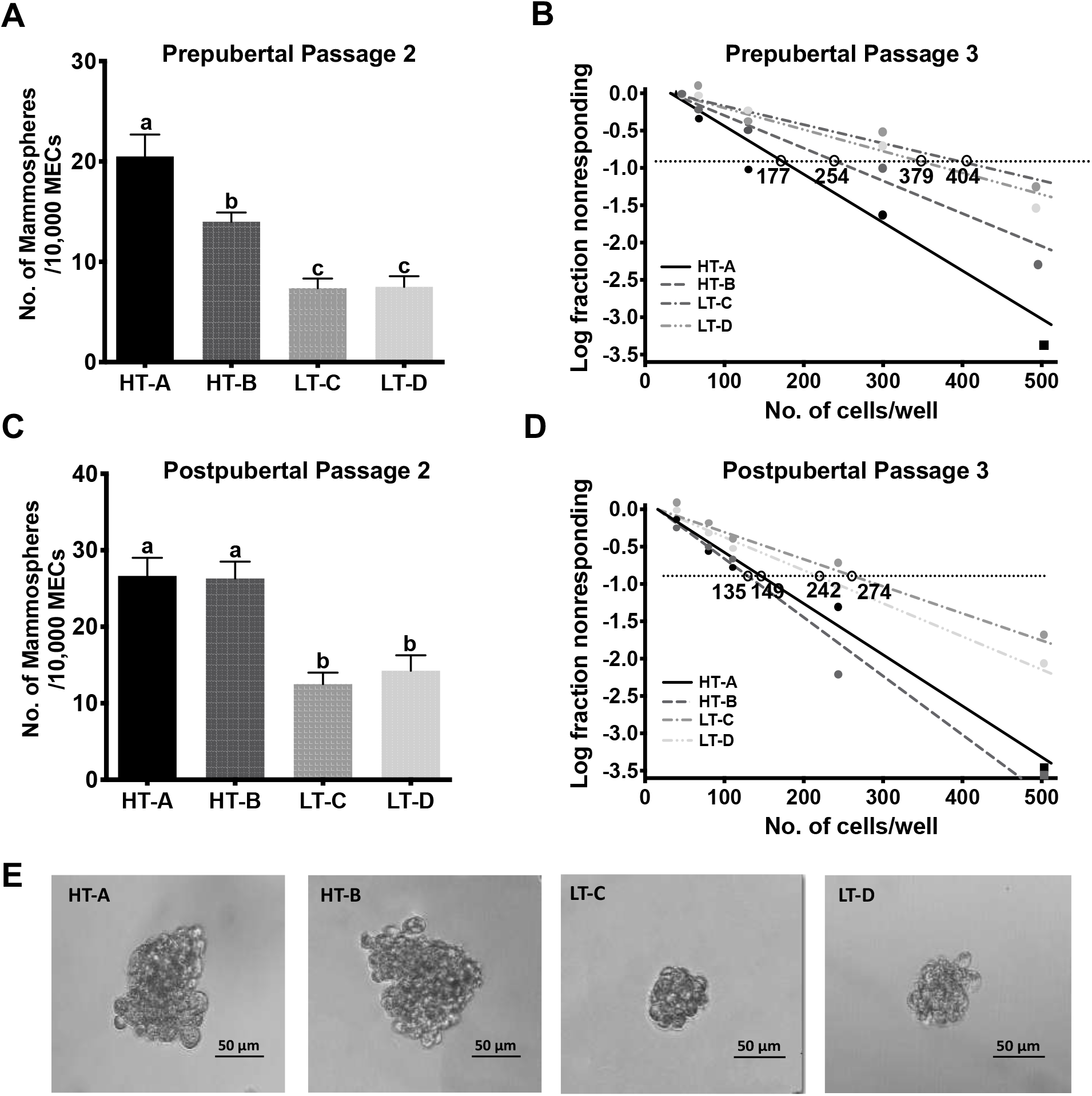
Mammosphere limiting dilution assays (LDAs) demonstrated HT group animals had a higher number of mammosphere-initiating cells than LT group animals. **(A)** Average numbers of mammospheres formed from prepubertal P2 MECs (n≥4). Mean ± SEM are shown. One-way ANOVA was used for statistical analysis and pairwise comparisons between groups were performed using Fisher’s LSD test. Columns are significantly different from each other if they do not share a letter (a≠b≠c, *P*<0.05). **(B)** LDA of prepubertal P3 mammospheres. The natural log fraction of non-responding wells plotted on a linear scale versus the cell density per well. **(C)** Average numbers of mammospheres formed from postpubertal HT and LT P2 MECs (n≥4). **(D)** LDA of postpubertal P3 mammospheres. The natural log fraction of non-responding wells plotted on a linear scale versus the cell density per well. **(E)** Representative images of postpubertal mammospheres from HT and LT groups.

In postpubertal animals, the numbers of P2 mammospheres formed by MECs from HT animals also increased by 2 times compared to those by either LT group (**Fig. 2C**). The LDA revealed that the mammosphere-forming efficiency of postpubertal cells from HT-A was 1.7-times higher compared to LT-C (1 in 149 versus 1 in 274, *P*<0.001) and LT-D (1 in 149 versus 1 in 242, *P*<0.005) (**Fig. 2D and Supplementary Table S2)**. Postpubertal HT-B animals also had a 1.9-fold increase in MaSC frequency when compared to LT-C (1 in 135 versus 1 in 274, *P*<0.001) and LT-D animals (1 in 135 versus 1 in 242, *P*<0.005) (**Fig. 2D**). Furthermore, we observed that the sizes of postpubertal mammospheres from both HT groups were larger (>100μM vs. 50-80μM in diameter) compared to both LT groups, indicating a higher proliferative capacity of progenitor cells that derived from HT group MaSCs (**Fig. 2E**). These data agree with the FACS results, indicating that the mammary tissue of HT animals harbored an increased number of highly clonogenic MaSCs compared to LT animals.

### IGF1 levels were increased and IGFBP5 levels were decreased in mammary tissues of HT animals compared to LT animals

We investigated which molecular mechanisms could drive this increase in the number of MaSCs in tissues from the HT groups compared to the LT groups. Human studies have shown that increased circulating IGF1 levels are associated with increased breast cancer risk [27], and in our previous study with this diet model, we found that serum IGF1 in 5-month old animals is significantly higher in the HT groups compared to the LT groups [3]. In addition, our studies in the BK5.IGF1 transgenic mouse model show that IGF1 overexpression in keratin 5-positive mammary basal epithelial cells increases the number of mammary terminal end buds, an early niche of MaSCs, within the developing mammary ducts [28,29]. Based on this evidence, we hypothesized that the HT dietary regimens increased the number of MaSCs through the upregulation of IGF1.

We first tested this hypothesis by measuring mammary tissue IGF1 levels in both pre- and postpubertal female mice. Prepubertal mammary glands from group HT-A expressed IGF1 mRNA levels that were 4-times higher than in both LT groups (**Fig. 3A**). IGF1 protein levels were also increased 2.6-times in group HT-A compared to the LT groups (**Fig. 3B**). Similarly, prepubertal HT animals from group HT-B expressed IGF1 mRNA levels that were 2.5-times higher than in the LT groups (**Fig. 3A**), and there was also a 1.6-fold corresponding increase in IGF1 protein levels in tissues from group HT-B compared to age-matched LT animals (**Fig. 3B**). In postpubertal glands, IGF1 mRNA and protein levels were also significantly higher in HT groups compared to LT groups (**Fig. 3A** and **B,** *P*<0.05). These results show that the HT dietary regimens significantly increased mammary tissue IGF1 levels during the prepubertal stage, and this increase persisted into adulthood.

**Figure 3.**
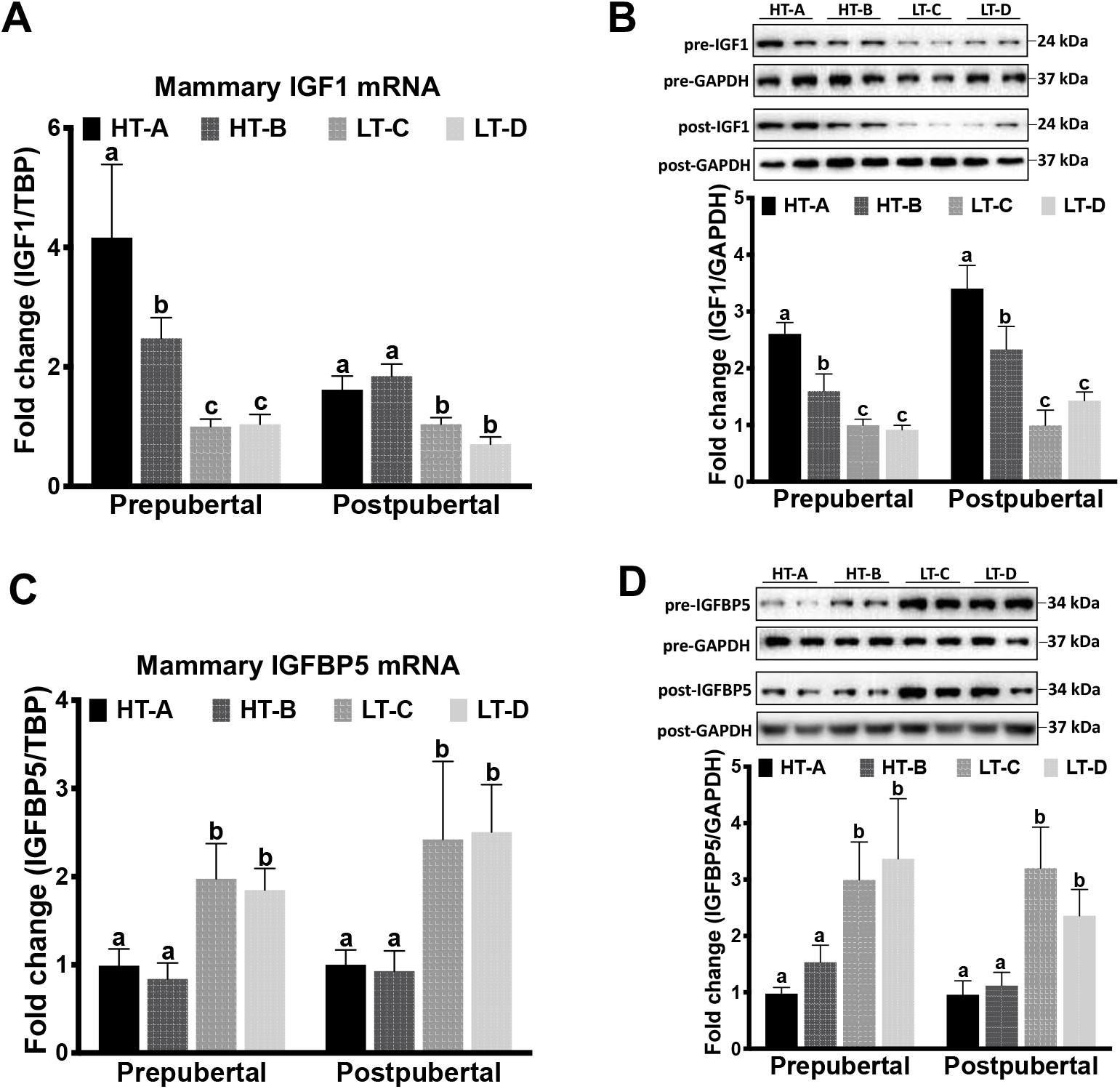
Mammary IGF1 levels increased in HT groups compared to LT groups, while IGFBP5 levels decreased. **(A)** Mammary tissue IGF1 mRNA levels of pre- and postpubertal animals from HT and LT groups (n≥4). RT-qPCR analyses are normalized to TATA-binding protein (TBP). Relative IGF1 mRNA levels are expressed as fold change compared to group LT-C. **(B)** IGF1 protein levels in pre- and postpubertal mammary tissues from HT and LT groups. Top panels, representative Western blots, bottom panels, densitometric quantifications (n≥6). **(C)** IGFBP5 mRNA levels in pre- and postpubertal mammary tissues from HT and LT groups (n≥4). Relative IGFBP5 mRNA levels are expressed as fold change compared to group HT-A**. (D)** IGFBP5 protein levels in pre- and postpubertal mammary tissues from HT and LT groups (n≥6). Mean ± SEM are shown. One-way ANOVA was used for statistical analysis, and pairwise comparisons were performed using Fisher’s LSD test. Groups are significantly different from each other if they do not share a letter (a≠b≠c, *P*<0.05).

The bioavailability of IGF1 in the mammary gland is regulated by IGF binding proteins (IGFBPs). *In vitro*, IGFBPs can inhibit IGF1 signaling by sequestering free IGFs from the IGF1 receptor (IGF1R) [30,31]. Six members of the IGFBP family (IGFBP1-6) are present in mammary tissue, of which IGFBP5 is the most prevalent [32]. We found that IGFBP5 mRNA levels were significantly lower in HT tissues compared to LT tissues at both the pre- and postpubertal stages (**Fig. 3C**, *P*<0.05). Likewise, prepubertal tissues from the HT groups harbored a 2.4-fold reduction in IGFBP5 protein levels compared to LT groups (**Fig. 3D**). Similarly, in tissues from postpubertal HT animals, IGFBP5 protein levels were also significantly lower than in tissues from LT animals (**Fig. 3D**, *P*<0.05). These findings indicate that the pro-tumorigenic dietary regimens of the HT animals not only increased total mammary IGF1 levels but also increased its bioavailability through the downregulation of IGFBP5.

### Increased IGF1 production in mammary stromal cells from HT animals promoted MaSC self-renewal

We next investigated whether the difference in tissue IGF1 levels between HT and LT animals affected the stemness and self-renewal capacity of MaSCs. Earlier studies demonstrate that IGF1 is highly expressed in mammary stromal cells and regulates mammary epithelial development in a paracrine fashion [33,34]. We therefore sorted the Lin^−^CD24^low^CD29^low^ mammary stromal cell population in prepubertal glands from HT and LT groups by FACS (Supplementary Fig. S1A). IGF1 mRNA levels in FACS-sorted mammary stromal cells were 5.6-times higher in group HT-A and 3.7-times higher in group HT-B compared to LT animals (**Fig. 4A**). In postpubertal stromal cells, IGF1 mRNA levels were also 2.9-times higher in HT groups compared to the LT groups.

**Figure 4.**
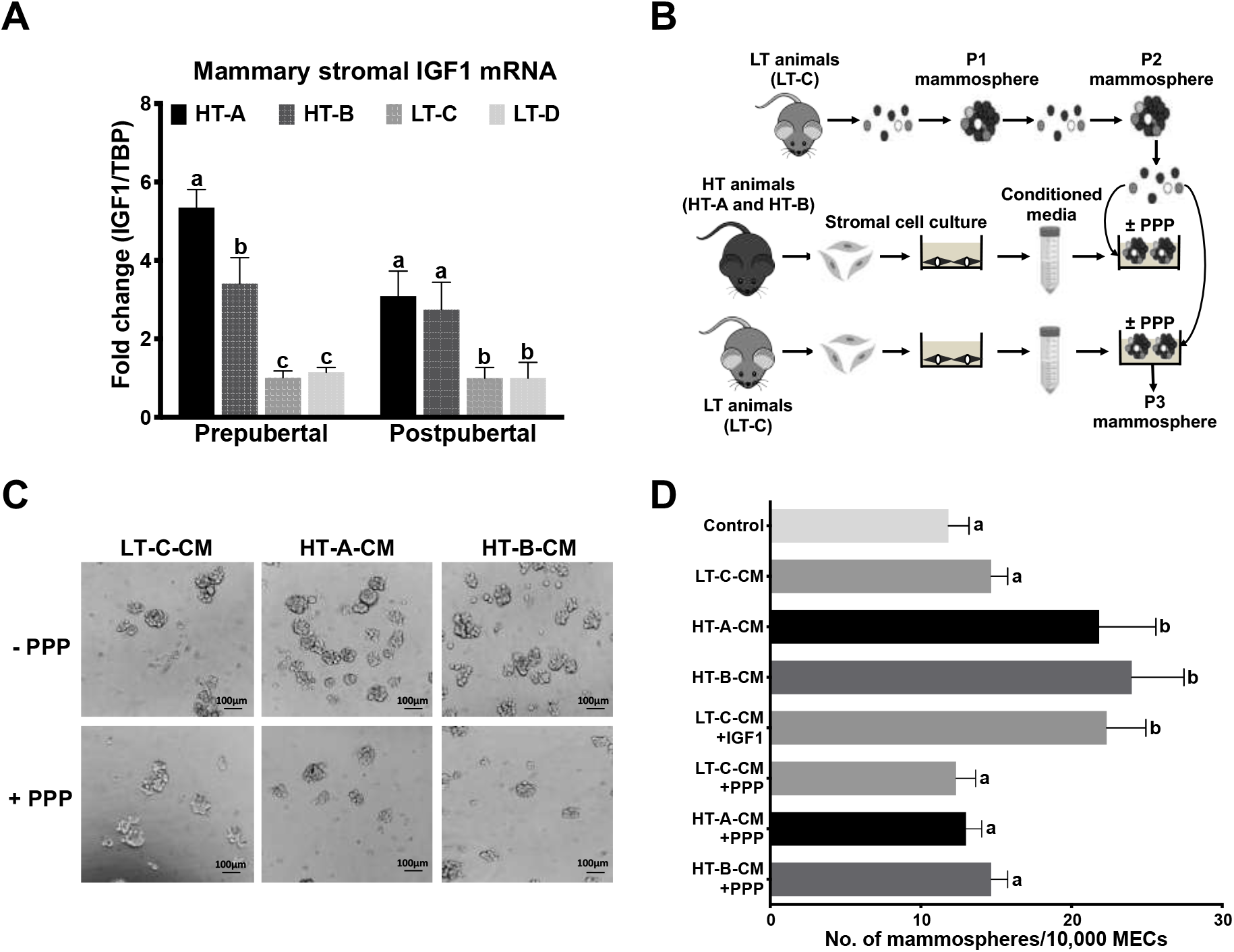
Increased IGF1 production in mammary stromal cells promoted MaSC self-renewal. **(A)** IGF1 mRNA in sorted mammary stromal cells from pre- and postpubertal tissues from HT and LT groups (n=4). Relative IGF1 mRNA levels are expressed as fold change from group LT-C**. (B)** Experimental procedure to perform the mammosphere assay using conditioned medium (CM) from stromal cell cultures. **(C)** Representative images of mammospheres formed in response to CM from mammary stromal cells from group LT-C (left), HT-A (center) and HT-B (right), ± picropodophyllin (PPP, lower panels). Scale bars indicated 100 μm. **(D)** Bar graph of numbers of mammospheres formed in CM from mammary stromal cells from group LT-C, HT-A and HT-B ±IGF1 or ±PPP (n=3). Mean ± SEM are shown. One-way ANOVA was used for statistical analysis. Pairwise comparisons were performed using Fisher’s LSD test. Groups are significantly different from each other if they do not share a letter (a≠b≠c, *P*<0.05).

To assess how this increased mammary stromal IGF1 expression affects the ability of MaSCs to self-renew, we performed a mammosphere-forming assay using medium conditioned by stromal cell cultures derived from either HT or LT group mammary tissues (**Fig. 4B**). Briefly, FACS-sorted mammary stromal cells from the prepubertal HT groups (HT-A and HT-B) and a LT group (LT-C) were plated separately with equal cell counts in serum-free stromal cell growth medium. Stromal cells were incubated for 24 hours and the supernatant (condition medium, CM) was collected. Prepubertal MECs from the group LT-C were harvested and cultured to generate P2 mammospheres as described previously. P2 mammospheres were then again disaggregated and re-plated using the previously harvested CM from either HT or LT mammary stromal cell cultures. Cells were then incubated in their respective CM for seven days to form P3 mammospheres.

As shown in **Fig. 4C** and **D**, CM produced by group LT-C (LT-C-CM) did not significantly affect mammosphere-forming efficiency compared to unconditioned mammosphere growth medium (control). Conversely, treating LT-C mammospheres with CM produced by group HT-A (HT-A-CM) or group HT-B (HT-B-CM), mammosphere numbers were significantly increased compared to those grown in LT-C-CM (**Fig. 4C** and **D**, *P*<0.04). These results indicated that mammary stromal cells from the HT groups secreted soluble factors that were able to promote the self-renewal capacity of MaSCs derived from a LT group.

To determine whether IGF1 was the relevant soluble factor that promoted MaSC self-renewal, we added recombinant IGF1 to LT-C-CM. Addition of recombinant IGF1 significantly increased the number of mammospheres formed in LT-C-CM, thus recapitulating the effect of HT-A-CM or HT-B-CM on promoting MaSC self-renewal (**Fig. 4D**, *P*<0.03). This result indicated that IGF1 was a plausible candidate factor. To confirm whether IGF1 was indeed the relevant soluble factor in our systems, we repeated the experiment by adding the IGF1 receptor (IGF1R) inhibitor picropodophyllin (PPP) to HT-A-CM, HT-B-CM and LT-C-CM. Adding PPP to LT-C-CM did not significantly affect the number of P3 mammospheres formed (**Fig. 4C** and **D**). However, when PPP was added to cultures grown in HT-A-CM or HT-B-CM, mammosphere numbers were significantly decreased compared to cultures grown in HT-A-CM or HT-B-CM without PPP (**Fig. 4C** and **D**, *P*≤0.01). Moreover, the numbers of mammospheres formed in HT-A-CM or HT-B-CM with addition of PPP were not different from those formed in LT-C-CM (**Figure 4D**). These results demonstrate that the increase in mammosphere numbers seen in HT-A-CM or HT-B-CM was due to the activation of IGF1R signaling. Together, these data show that the pro-tumorigenic dietary regimens increased IGF1 production in mammary stromal cells, which then promoted MaSC self-renewal in a paracrine fashion.

### The pro-tumorigenic dietary regimens decreased mammary stromal DNMT3b levels and induced DNA hypomethylation of the IGF1 promoter 1

Early environmental exposures can affect gene transcription through epigenetic alterations, such as DNA methylation (reviewed in [35]). DNA methylation of the dinucleotide CG represses gene expression through direct or indirect inhibition of transcription factor binding to the gene promoters [36]. We next tested whether the increase in IGF1 transcription in mammary stromal cells from HT animals resulted from DNA hypomethylation of the IGF1 promoters.

IGF1 transcription is driven by two alternative promoters (**Fig. 5A**). Promoter 1 (Pr1) initiates transcription from exon 1 and gives rise to the Class 1 transcript, while promoter 2 (Pr2) initiates from exon 2, producing the Class 2 transcript. In rodents, Pr1 becomes active during the embryonic stage and remains active through adulthood, whereas Pr2 remains silent until 3 weeks of age when growth hormone exerts its effect by stimulating IGF1 expression [37]. Both transcript isoforms generate the same mature IGF1 peptide [38]. QPCR analysis revealed that both Class 1 and Class 2 transcripts were expressed at significantly higher levels in FACS-sorted mammary stromal cells from prepubertal HT animals compared to LT animals (*P*<0.05) (**Fig. 5B**). We then used bisulfite sequencing analysis to investigate whether this difference in Class 1 and Class 2 mRNA levels was induced by differential DNA methylation of CG sites that flank the transcription start sites (TSS) of the Pr1 and Pr2 promoters. As shown in **Fig. 5C** and **D**, in mammary stromal cells from prepubertal animals, the Pr1 in both HT groups was hypomethylated compared to the LT groups on four out of five CG sites that flank the TSS: 41.0% of the CG-239 sites were methylated in the HT groups compared to 72.0% in the LT groups (*P*<0.05); CG-142, 15.5% in HT groups vs. 48.5% in LT groups (*P*<0.01); CG-110, 27.5% in HT groups vs. 44.5% in LT groups (*P*<0.08); CG-78, 16.5% in HT groups vs. 35% in LT groups (*P*<0.01). The combined (total) methylation percentage of these five CG sites was also significantly decreased in mammary stromal cells from the HT groups compared to the LT groups (**Fig. 5D**, *P*<0.01). In addition, the amounts of IGF1 Class 1 mRNA in these tested samples were reversely correlated with the combined DNA methylation percentage of the five CG sites (**Fig. 5E**, R^2^=0.77, *P*<0.0001). These results indicate that the pro-tumorigenic dietary regimens increased IGF1 expression in mammary stromal cells from HT animals by inducing DNA hypomethylation of the Pr1 promoter.

**Figure 5.**
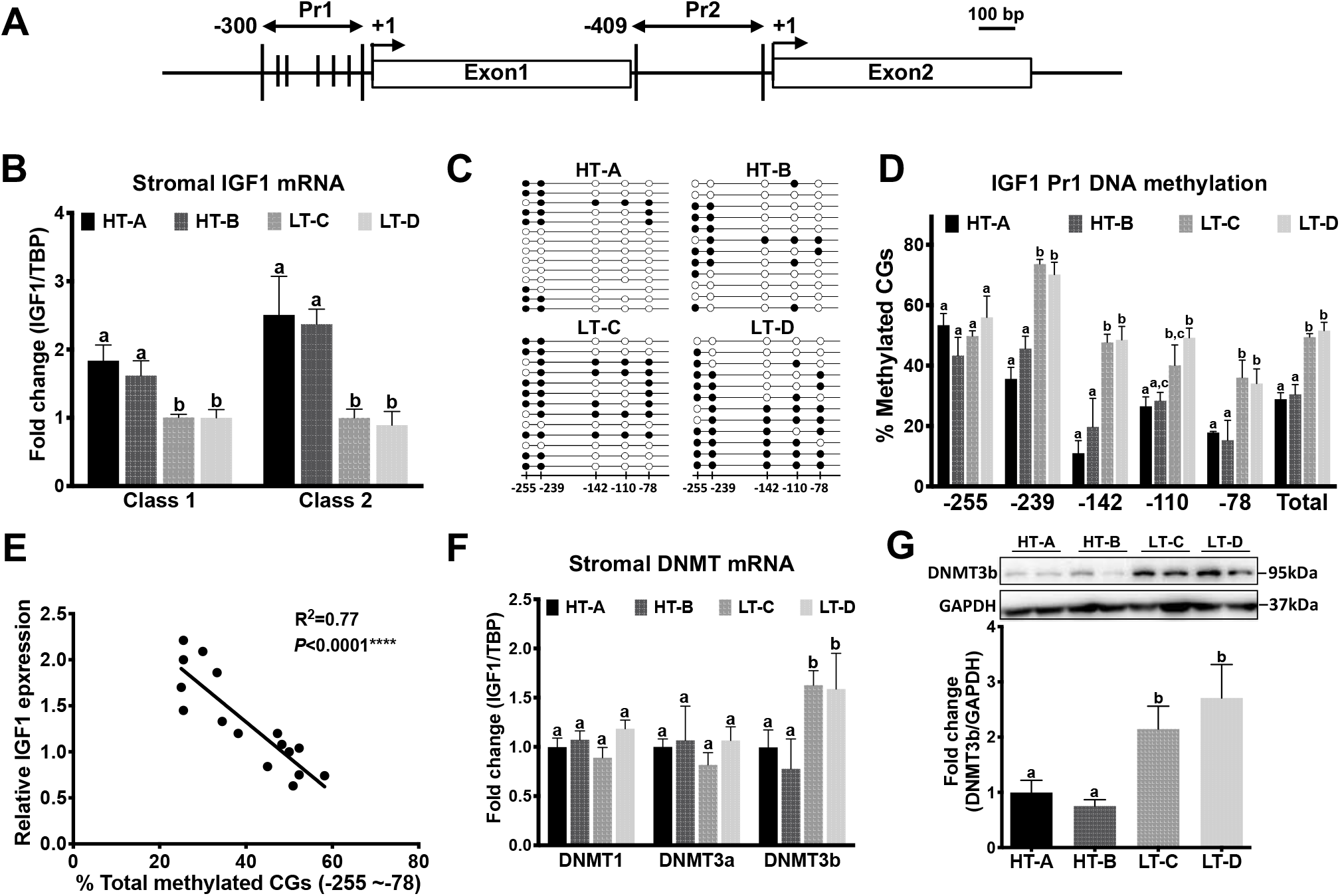
DNA methylation of the IGF1 Pr1 promoter was decreased in mammary stromal cells from prepubertal HT animals and DNMT3b levels were also decreased. **(A)** Graphical representation of the mouse IGF1 promoters 1 (Pr1) and 2 (Pr2). Transcription start sites (TSSs, +1) are shown as arrows, CG sites used for the analysis shown as vertical bars. **(B)** mRNA levels of Class 1 and Class 2 transcripts in mammary stromal cells from HT and LT groups (n=4). RT-qPCR analyses were normalized to TBP. Relative mRNA levels are expressed as fold change from group LT-C. **(C)** Representative DNA methylation patterns on the Pr1 from one HT and one LT animal from each group. Each circle represents a CG site as shown in panel (A), each line represents a single clone, and closed circles show methylated CG sites. **(D)** Quantification of DNA methylation percentages for each CG site (n≥4). **(E)** Correlation scatter plot with best fit line representing the Class 1 transcript mRNA levels with the combined methylation percentage of the five CG sites at IGF1 Pr1 (Pearson’s correlation test, R^2^=0.77, *P*<0.0001, one-tailed). **(F)** mRNA levels of DNA methyltransferases (DNMTs) in sorted mammary stromal cells from postpubertal tissues (n=4). RT-qPCR analyses were normalized to TBP. Relative mRNA levels are expressed as fold change from group D. (**G**) DNMT3b protein levels in stromal cells from HT and LT groups (n≥4). Representative Western blot images (top panel) and densitometric quantifications (bottom panel). Mean ± SEM are shown. One-way ANOVA was used for statistical analysis. Pairwise comparisons were performed using Fisher’s LSD test. Groups are significantly different from each other if they do not share a letter (a≠b≠c, *P*<0.05).

Since the Class 2 transcript was also significantly upregulated in HT animals (**Fig. 5B**), we also analyzed DNA methylation levels on six CG sites (CG-371-CG-255) that flank the Pr2 TSS (Supplementary Fig. S2A). However, we found that these CG sites were largely hypomethylated in both HT and LT groups, suggesting that DNA methylation in other genomic regions, such as distal enhancers, or other epigenetic mechanisms, such as histone modifications, may contribute to the differential expression of the Class 2 transcript in these animals (Supplementary Fig. S2B and C).

Genomic DNA methylation is catalyzed by DNA methyltransferases (DNMTs), including DNMT1, DNMT3a and DNMT3b. Therefore, we measured gene expression levels of these DNMTs in FACS-sorted stromal cells from prepubertal mammary tissues. We found that mammary stromal cells from HT groups had significantly lower DNMT3b mRNA levels compared to the LT groups (*P*<0.05), whereas gene expression levels of DNMT1 and DNMT3a were not different (**Fig. 5F**). Western blot analysis showed that DNMT3b protein levels were also significantly decreased in mammary stromal cells from HT groups compared to those from LT groups (**Fig. 5G,** *P*<0.03). These results suggest that the pro-tumorigenic DR/ _ /HF dietary regimens induced IGF1 promoter hypomethylation that is mediated by decreased DNMT3b levels.

We then investigated a potential causal relationship between DNA hypomethylation of the IGF1 Pr1 and decreased DNMT3b levels in murine cells. Since fibroblasts are one of the predominant cell types in mammary stroma [39], we knocked down DNMT3b by transfecting DNMT3b siRNA into NIH 3T3 cells, a mouse embryonic fibroblast cell line. DNMT3b siRNA transfection reduced DNMT3b mRNA levels by 60% compared to scrambled controls 48 hours post-transfection (**Fig. 6A**). In contrast, mRNA levels of DNMT1 and DNMT3a were not significantly changed, indicating a high specificity of the DNMT3b knockdown (**Fig. 6A**). DNMT3b protein levels were also decreased by 65% in siRNA-treated cells compared to controls (**Fig. 6B**). To assess whether this DNMT3b knockdown resulted in IGF1 Pr1 hypomethylation, we performed bisulfite sequencing analyses of IGF1 Pr1 in these DNMT3b siRNA-transfected cells. As shown in **Fig. 6C**, DNMT3b knockdown significantly decreased IGF1 Pr1 methylation of all five CG sites (−255 ~ −78) that flank the TSS (*P*<0.01). Concordant with this CG hypomethylation, IGF1 mRNA levels were significantly increased in DNMT3b siRNA-transfected cells compared to the controls (**Fig. 6D**). These results show that DNMT3b knockdown induced IGF1 Pr1 hypomethylation, which then resulted in an increase in IGF1 mRNA levels.

**Figure 6.**
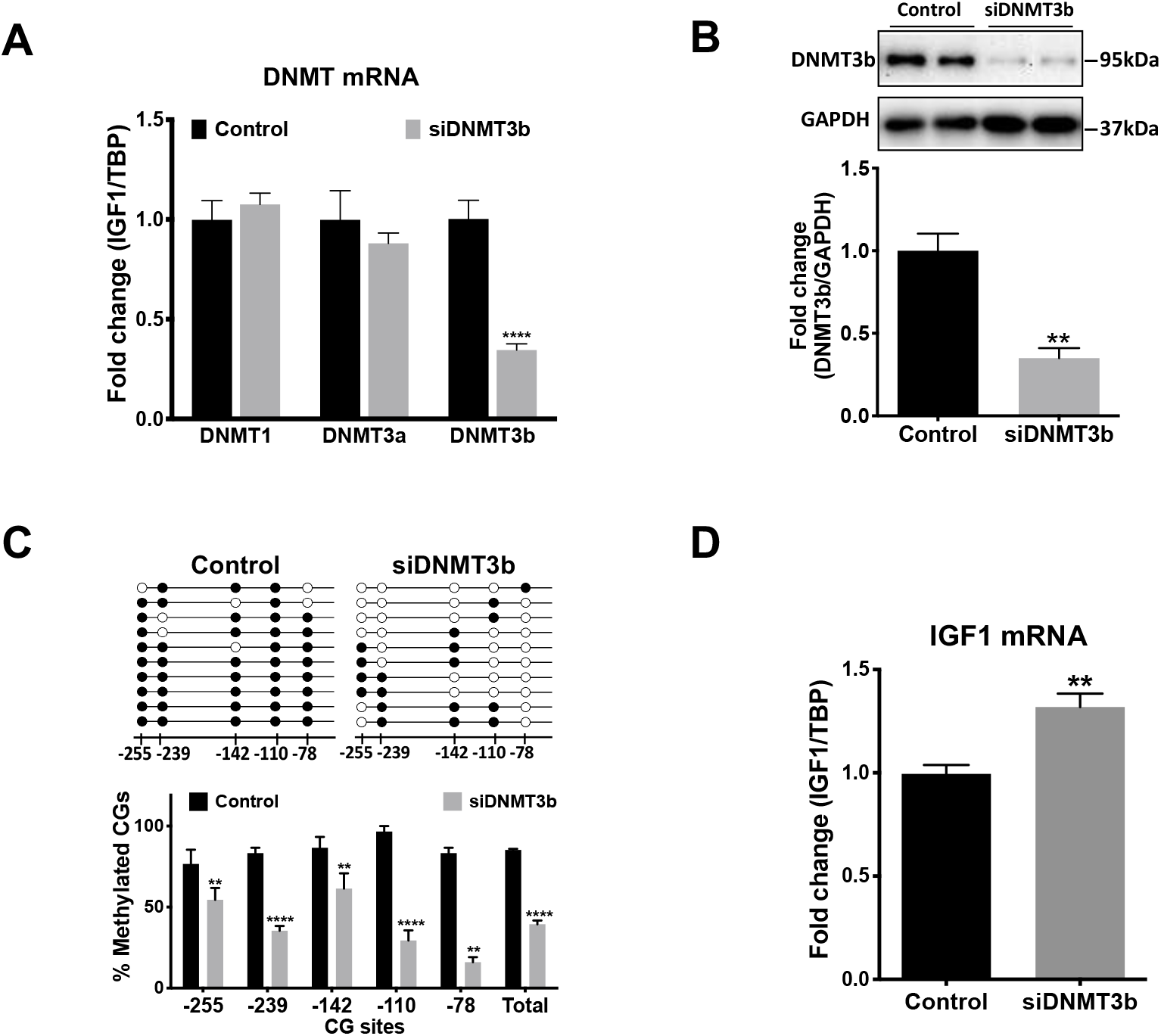
DNMT3b knockdown decreased IGF Pr1 DNA methylation and increased IGF1 mRNA expression. **(A)** mRNA levels of DNMTs in 3T3 cells transfected with scrambled siRNA (control) or DNMT3b siRNA (siDNMT3b). RT-qPCR analyses were normalized to TBP (n=4). **(B)** DNMT3b protein levels in control and siRNA treated groups. Representative Western blot images (top panel) and densitometric quantifications (bottom panel, n=4). **(C)** Representative DNA methylation patterns of the IGF1 Pr1 (top panel) and quantifications of methylated CG sites (bottom panel, n=3). **(D)** Total mRNA levels of IGF1 (n=4). Mean ± SEM are shown. Pairwise comparisons were performed using Student’s t-test. \*\**P*<0.01, \*\*\**P*<0.001, \*\*\*\**P*<0.0001.

### The pro-tumorigenic dietary regimens promoted the formation of adenosquamous carcinomas

Previous findings have demonstrated that tumors initiating from different mammary epithelial cell types develop into distinct histopathologic phenotypes [13,40,41]. Keller *et al*. reported that Ras-transformed human breast basal epithelial cells produce metaplastic tumors with squamous differentiation, and that these tumors highly express the basal lineage marker keratin 14 (K14). In contrast, Ras-transformed luminal cells produce ductal adenocarcinomas with predominantly luminal features, including protein expression of the luminal lineage markers K8 and K19 [41]. Their transcriptomic analyses show that metaplastic tumors derived from RAS-transformed breast basal epithelial cells highly express mRNA of previously identified MaSC signature genes [41]. Since we observed that tissues from HT animals had an increased proportion of MaSCs compared to LT animals, we tested whether this difference was associated with development of distinct tumor phenotypes in HT and LT groups. Therefore, we reanalyzed histological sections of mammary tumors that were collected from our previous carcinogenesis study [3]. Fourteen tumors from the HT groups (seven from HT-A and seven from HT-B) and eleven tumors from the LT groups (seven from LT-C and four from LT-D) were analyzed (Supplementary Table S3). In the HT groups, seven tumors (50%) were classified as metaplastic carcinomas of either squamous or adenosquamous phenotype, which show noticeable squamous metaplasia and widespread keratin pearls (**Fig. 7A** and **B**). Seven tumors (50%) were classified as ductal adenocarcinomas of either acinar, papillary or solid type, typically with central necrosis (Supplementary Table S3). Conversely, in the LT groups, only one tumor (0.9%) was classified as an adenosquamous carcinoma; the majority of tumors (99.1%) were ductal adenocarcinomas of acinar, papillary or solid phenotype (**Fig. 7A** and **B**). The overall proportion of squamous or adenosquamous (metaplastic) carcinomas was significantly higher in HT groups than in LT groups (**Fig. 7B**, Chi-square test, *P*=0.03). Immunohistochemistry (IHC) showed that adenosquamous carcinomas from the HT groups had significantly higher expression of the mammary basal lineage marker K5 and low expression of the luminal lineage marker K8 compared to the ductal adenocarcinomas from the LT groups (**Fig. 7C** and **D**, *P*<0.01). These results indicate that the adenosquamous carcinomas from the HT groups may have derived from cells of the basal/stem cell compartments, suggesting that early exposure to the pro-tumorigenic DR/_ /HF diet promoted the formation of adenosquamous tumors in HT animals by expanding the MaSC population.

**Figure 7.**
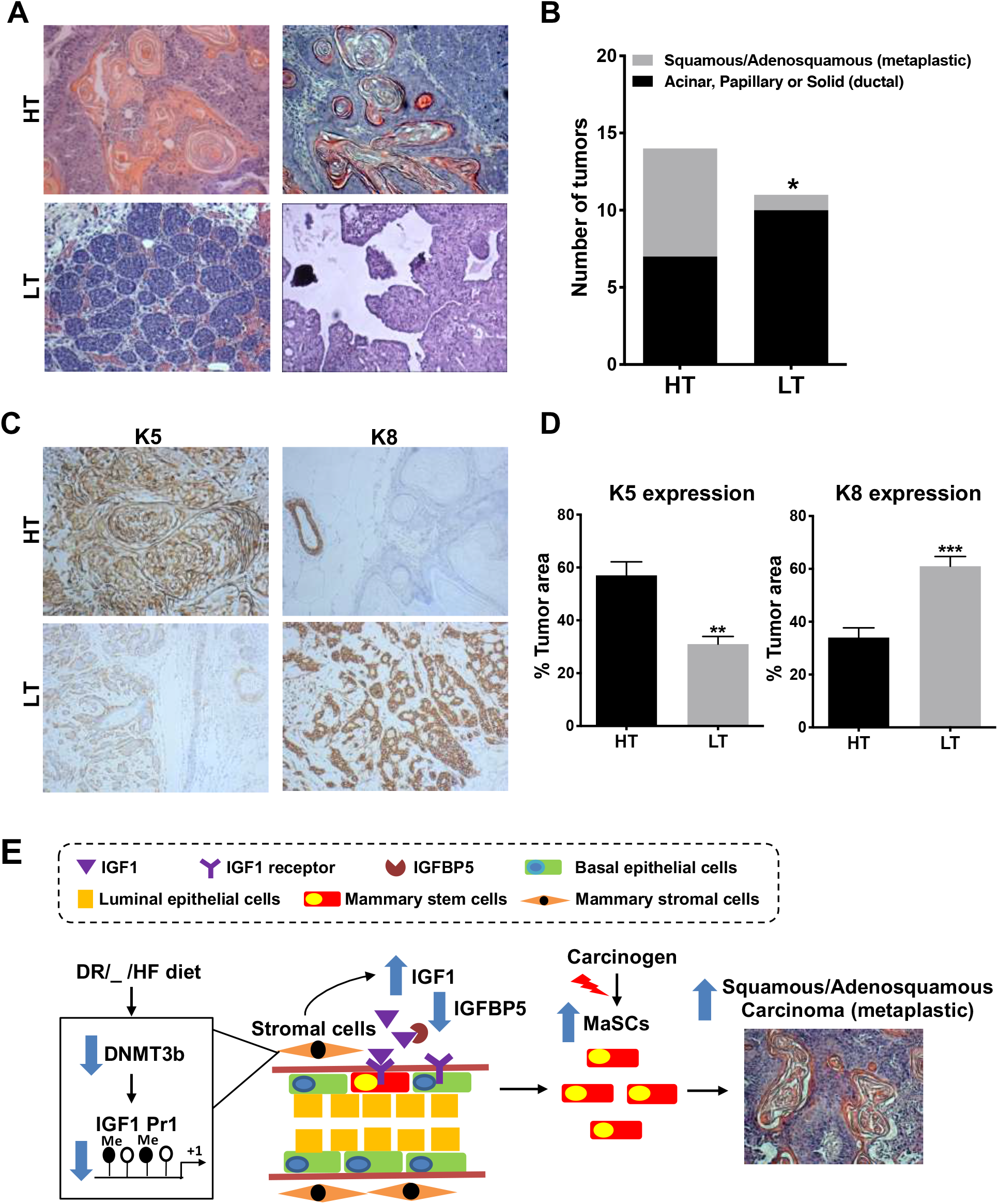
HT and LT groups showed differences in proportions of specific histopathologic tumors types. (**A**) Representative hematoxylin and eosin (H&E) stained tumors from HT groups and LT groups. Top left, a squamous tumor from a HT-B animal. Top right, an adenosquamous tumor from a HT-A animal. Bottom left, a ductal adenocarcinoma of acinar type from a LT-C animal. Bottom right, a ductal adenocarcinoma of papillary type from a LT-D animal. Pictures were taken at 10X magnification. **(B)** Prevalence of histopathological phenotypes of tumors arising in the HT groups and LT groups. Tumor prevalence was analyzed by chi-squared (χ^2^) test. (**C**) IHC staining of K5 (left panels) and K8 (right panels) for tumors developed from a HT-B animal (top panels) and a LT-C animal (bottom panels). **(D)** Quantification of K5 and K8 protein expression in tumors developed from HT and LT groups animals (n=7 per group). Student’s t-test was used for statistical analysis. **(E)** Proposed model of the effect of the pro-tumorigenic dietary regimens of HT animals on MaSCs.

## Discussion

To propose effective cancer prevention strategies, it is critical to identify molecular and cellular mechanisms through which environmental exposures program disease susceptibilities at early developmental stages. In this study, we demonstrated that early dietary manipulations during gestation, lactation and postweaning programmed lifetime mammary cancer risk by increasing the number of carcinogen-susceptible MaSCs through sustained activation of the IGF1 signaling pathway into adulthood. Based on the data presented, we propose the following model (**Fig. 7E**): The pro-tumorigenic DR/_ /HF dietary regimens induced DNA hypomethylation of the IGF1 Pr1 promoter in mammary stromal cells of HT animals through decreased DNMT3b levels, resulting in increased stromal IGF1 expression. These pro-tumorigenic dietary regimens also decreased IGFBP5 levels in mammary tissues from the HT animals, resulting in increased bioavailability of IGF1. Increased IGF1 production and bioavailability promoted MaSC self-renewal and expanded the MaSC population before the animals reached puberty, thereby increasing the number of carcinogen-susceptible targets later in life.

Mammary stem and progenitor cells are potential tumor-initiating cells [12–14], and the lifetime risk for developing breast cancer is partly determined by the number of these cells in the mammary gland [3,14–16]. Our flow cytometric data show that HT animals had a significantly increased proportion of MECs with MaSC surface markers than LT animals and this difference was observed before animals reached puberty (**Fig. 1**). Concordant with this, mammosphere limiting dilution assays show that MECs from the HT animals formed significantly more secondary and tertiary mammospheres than the LT animals (**Fig. 2**). These results demonstrated that the pro-tumorigenic DR/ _ /HF dietary regimens increased the size of the MaSC compartment in HT animals before puberty. Since prepubertal animals from group HT-B had been exposed to the postnatal HF diet for only two weeks, from 3 to 5 weeks of age, we reason that the expansion of the MaSC population in HT animals is independent from developing obesity. These results indicate how mammary tumor susceptibility can be programmed during the early stages of mammary gland development, demonstrating that successful breast cancer prevention strategies should be applied in humans starting in infancy and during childhood.

Early dietary exposures have been shown to program lifetime disease susceptibilities through epigenetic mechanisms [42]. Epigenetic regulation of IGF1 gene expression has been reported in the liver, hematopoietic systems, and placenta [43–45]. *Fung et al*. showed that intrauterine growth restriction induced by maternal vasoconstriction increases DNA methylation at a specific CG site (−78) of the IGF1 Pr1 and represses IGF1 transcription in the liver of juvenile male mice [45]. Ouni *et al.* reported that DNA methylation of the IGF1 Pr2 in mononuclear cells of peripheral blood is negatively associated with serum IGF1 levels and body stature in a Caucasian children cohort [46]. These results indicate that IGF1 expression is epigenetically regulated through DNA methylation at the Pr1 and Pr2 promoters in both rodents and humans, and that these epigenetic alterations induce specific biological outcomes. In this study, we present for the first time that switching from a healthy DR diet to a HF diet reprogrammed IGF1 gene expression in mammary tissue by inducing DNA hypomethylation of the Pr1 promoter, and that this DNA hypomethylation was mediated by decreased mRNA and protein levels of DNMT3b in mammary stromal cells. When comparing tissue-specific differentially methylated regions (DMRs) of the genomes between humans, mice and rats, Zhou *et al*. reported that up to 37% of the DMRs are conserved between humans and rodents, and this epigenetic conservation is largely driven by the similarity of the primary genomic DNA sequence between species [47]. The nucleotide sequences of IGF1 promoters are highly conserved between humans and rodents (85%-92% sequence similarity) [48]. Therefore, the diet-induced epigenetic alteration of the IGF1 Pr1 in mice likely recapitulates a similar effect in humans consuming a Western diet starting in infancy and during childhood. Since this epigenetic alteration of the IGF1 Pr1 was observed in prepubertal mammary stromal cells, and DNA methylation is a stable epigenetic marker that can be inherited through multiple cell divisions [49], our results further indicate that early, prepubertal dietary intervention is a critical component of successful breast cancer prevention strategies.

Breast cancer can be divided into different subtypes based on histopathologic phenotypes and gene expression profiles, leading to different treatment strategies and outcomes (reviewed in [50]). We show that early exposure to the pro-tumorigenic DR/ _ /HF dietary regimens promoted the formation of mammary metaplastic carcinomas of either the squamous or adenosquamous phenotype in carcinogen-treated mice. Mammary adenosquamous carcinomas are shown to have shorter latencies compared to ductal adenocarcinomas in studies of carcinogen-treated mice or transgenic mice with different genetic background [28,51,52], and adenosquamous carcinomas are of triple-negative (ER^−^, PR^−^ and HER2^−^) phenotype in both rodents and humans [51,53]. Comparatively, carcinogen-induced ductal adenocarcinomas of either acinar, papillary or glandular phenotype are hormone-receptor positive (ER^+^, PR^+^), suggesting that these tumors may have a better prognosis and treatment outcomes than adenosquamous carcinomas [51]. These results indicate that early dietary manipulations affect tumor incidence and program tumor phenotypes, which will ultimately affect cancer treatment strategies and outcomes.

Evidence from lineage tracing and transcriptome studies indicates that mammary tumors of different histopathological phenotypes are derived from different mammary epithelial cell types. Hollern *et al.* reported that mammary squamous carcionomas from MMTV-PyMT transgenic mice highly express previously identified gene signatures of MaSCs, whereas tumors of ductal glandular phenotypes highly express gene signatures of luminal progenitors or mature luminal cells [54]. Concordant with this, Keller *et al.* reported that metaplastic tumors that are derived from Ras-transformed human EpCAM^low^CD10^+^ breast basal epithelial cells highly express gene signatures of MaSCs compared to ductal adenocarcinomas that are derived from EpCAM^high^CD10^−^ luminal cells [55]. These results indicate that metaplastic carcinomas within mammary tissue may have derived from MaSCs, which would agree with our data showing that HT animals had an expanded MaSC population and developed increased proportions of adenosquamous carcinomas compared to LT animals. These results suggest that early dietary manipulations may result in different susceptibilities to specific mammary tumor phenotypes by affecting homeostasis of the mammary epithelial cell population.

In summary, our results show that the pro-tumorigenic DR/ _ /HF dietary regimens programmed lifetime mammary cancer susceptibility before puberty, and this programming effect was mediated by DNA hypomethylation of the IGF1 Pr1 in mammary stromal cells. This epigenetic alteration increased IGF1 expression in mammary tissues and expanded the MaSC compartment before puberty, resulting in a higher number of susceptible carcinogen targets. We demonstrated that mammary cancer susceptibility can be programmed before puberty, showing that dietary intervention at an early developmental stage is an essential component of successful prevention strategies.

## Materials and Methods

### Mice and Diets

All experimental animal procedures were approved by the Institutional Animal Care and Use Committee (IACUC). Animal maintenance and dietary treatment were performed as described previously [3]. Briefly, SENCAR breeder mice [56] were maintained at our animal facility under a 12-hour light/dark cycle at 24°C. Defined diets were purchased from Research Diets, Inc. Cat#D01060501 was the chow-like, low fat, low sugar control diet, and cat#D04011601, the high fat, high sugar (HF) diet. Female breeder mice were separated into control and HF diet groups at four weeks of age. As these animals were sedentary and commonly became obese with age [3], a mild 12% diet restriction was imposed on the control diet group (DR), except during lactation, which ensures proper nutrition of mothers and pups. This mild portion control did not result in an insufficiency of macro- or micronutrients. HF animals were also provided with 10% fructose in their drinking water.

HF mice became glucose intolerant after ten weeks compared to DR controls, as determined by glucose tolerance tests [3]. At 15 weeks of age, normal DR and hyperglycemic HF mothers were mated with normal SENCAR males. Upon birth, pups were fostered within 24 hours of birth to separate into the lactation exposure groups. At weaning, pups were randomized into the different postweaning diet exposure groups and were maintained on that diet until being euthanized.

### Mammary Epithelial Cell Isolation

Details for mammary epithelial cell (MEC) isolation are described in the Supplementary Materials and Methods. Fourth and fifth mammary gland pairs were resected from mice and were enzymatically digested with collagenase/hyaluronidase (Stem Cell Technologies, cat #07919) following the manufacturer’s protocol. Cells were then treated with 2ml of 0.25% Trypsin/EDTA solution (Stem Cell Technologies, cat #07901) and 2ml of 5mg/mL dispase (Gibco, cat#17105-041).

### Flow Cytometry and Cell Sorting

Details for flow cytometry and cell sorting are described in the Supplementary Materials and Methods. Antibodies and dilutions used for cell staining were shown in the Supplementary Table S4. To analyze epithelial populations, stained cells were loaded onto a Fortessa X20 flow cytometer (BD Biosciences). To sort mammary stromal cells, a FACSAria II flow cytometer (BD Biosciences) was used. Sorted cell populations were reanalyzed and found to be 94%-98% pure, and cell viability was above 85%.

### Mammosphere Culture and Limiting Dilution Assay

Details for mammosphere culture, limiting dilution assays and media components are described in the Supplementary Materials and Methods. MECs were plated at 10,000 cells/well in pHEMA-coated 24-well plates and cultured for seven days on P1 and P2 passages. Single cells from P2 mammospheres were plated into a 96-well ultra-low attachment plate at dilutions ranging from 1 to 512 cells per well (8 replicates per dilution). After seven days, wells were scored for the presence or absence of P3 mammospheres. To assess the frequency of mammosphere-initiating cells, an extreme limiting dilution analysis was performed as described previously [26]. Pairwise differences between the groups were compared with likelihood ratio tests using the asymptotic chi-squared (χ2) test approximation to the log–ratio [26].

For performing CM assays, single cells from P2 mammospheres were plated at 10,000 cells/well in pHEMA-coated 24-well plates with the respective CM used in the study. Recombinant IGF1 (Life Technologies, cat# PHG0071) and picropodophyllin (PPP, Santa Cruz, cat# sc-204008) were added to the CM at a final concentration of 7.5 nM and 10 μM, respectively.

### RT-qPCR and Western Blot Analysis

RT-qPCR and Western blot analysis were performed as described in the Supplementary Materials and Methods. Primer sequences for RT-qPCR are listed in the Supplementary Table S5. Target gene expression was normalized to TATA-binding protein (TBP). Antibodies and dilutions for Western blot analysis were listed in the Supplementary Table S4. Signals of target protein bands were normalized to GAPDH bands of the same sample and then normalized to the control group to calculate fold changes.

### DNA Methylation Analysis

Genomic DNA was extracted from mammary stromal cells using the DNeasy Blood &Tissue kit (Qiagen Cat #69504). Bisulfite conversion of genomic DNA was performed with a sodium bisulfite kit (EZ DNA Methylation-Lightning Kit, Cat #D5030) following the manufacturer’s instructions. Bisulfite primers used to cover CG sites on Pr1 and Pr2 are listed in Supplementary Table S5. PCR conditions were 95°C for 3min, 95°C for 15s, annealing at 55°C for 20s, 72°C for 20s, 38 cycles. PCR products were cloned into the T-vector pMD19 (TAKARA Bio Inc, Cat#3271). Plasmid DNA was extracted and sequenced with M13 forward or reverse primers (Eton Bioscience Inc, San Diego, CA). DNA methylation analysis was performed using the Quantification tool for Methylation Analysis (QUMA, http://quma.cdb.riken.jp/top/index.html) [57].

### siRNA Transfection

NIH 3T3 cells were grown to 70% confluency in DMEM/Ham’s F-12 (Caisson Labs, cat#DFP17) culture media with addition of 10% FBS. Cells were trypsinized and replated in 12-well plates. After reaching 50% confluency, cells were treated with Lipofectamine RNAiMAX Transfection Reagent (Invitrogen, cat #13778100) with addition of 20 nM DNMT3b siRNA (ThermoFisher, cat #161533) according to manufacture protocols. Cells were then maintained in DMEM/Ham’s F-12 culture media with addition of 2% FBS for 48 hours before collection.

### Histological Analysis

Preparations of histological sections and H&E or IHC staining were performed by the histology laboratory of Texas A&M University. Mammary tumor subtypes were classified referring to the classification systems proposed by Cardiff *et al.* [58].

### Statistical Analysis

Numerical results reflect mean ± SEM. Comparisons between groups were performed using one-way ANOVA and pairwise comparisons were conducted using Fisher’s LSD test unless otherwise specified. Statistical analyses were conducted, and graphical representations of data were plotted using the GraphPad Prism 6 software. *P* < 0.05 was considered significant.

## Acknowledgements

We appreciate Dr. Sarah Bondos, Dr. Kayla Bayless and Dr. Carl Gregory in the Department of Molecular and Cellular Medicine, Texas A&M University for their careful review of this manuscript. We thank Dr. Huijuan Yan (University of California, San Francisco) for her advice in designing and creating the figures in this manuscript.

## Notes

**Conflict of Interest Statement:** The authors declare no potential conflicts of interest.

### Competing Interest Statement

The authors have declared no competing interest.

## References

[1] Govindarajah V, Leung YK, Ying J, Gear R, Bornschein RL, Medvedovic M, et al. In utero exposure of rats to high-fat diets perturbs gene expression profiles and cancer susceptibility of prepubertal mammary glands. J Nutr Biochem 2016;29:73–82.

[2] ainter RC, De Rooij SR, Bossuyt PMM, Osmond C, Barker DJP, Bleker OP, et al. A possible link between prenatal exposure to famine and breast cancer: A preliminary study. Am J Hum Biol 2006;18:853–6.

[3] Lambertz IU, Luo L, Berton TR, Schwartz SL, Hursting SD, Conti CJ, et al. Early Exposure to a High Fat/High Sugar Diet Increases the Mammary Stem Cell Compartment and Mammary Tumor Risk in Female Mice. Cancer Prev Res 2017;10:553–62.

[4] Elias SG, Peeters PHM, Grobbee DE, van Noord PAH. Breast cancer risk after caloric restriction during the 1944-1945 Dutch famine. J Natl Cancer Inst 2004.

[5] de Oliveira Andrade F, Fontelles CC, Rosim MP, de Oliveira TF, de Melo Loureiro AP, Mancini-Filho J, et al. Exposure to lard-based high-fat diet during fetal and lactation periods modifies breast cancer susceptibility in adulthood in rats. J Nutr Biochem 2014;25:613–22.

[6] de Assis S, Warri A, Cruz MI, Laja O, Tian Y, Zhang B, et al. High-fat or ethinyl-oestradiol intake during pregnancy increases mammary cancer risk in several generations of offspring. Nat Commun 2012;3:1053.

[7] Hilakivi-Clarke L, Clarke R, Onojafe I, Raygada M, Cho E, Lippman M. A maternal diet high in n - 6 polyunsaturated fats alters mammary gland development, puberty onset, and breast cancer risk among female rat offspring. Proc Natl Acad Sci U S A 1997;94:9372–7.

[8] Semir Beyaz, Mana MD, Roper J, Kedrin D, Saadatpour A, Hong S-J, et al. High-fat diet enhances stemness and tumorigenicity of intestinal progenitors. Nature 2016;531:53–8.

[9] DeClercq V, McMurray DN, Chapkin RS. Obesity promotes colonic stem cell expansion during cancer initiation. Cancer Lett 2015;369:336–43.

[10] Shackleton M, Simpson KJ, Stingl J, Smyth GK, Wu L, Lindeman GJ, et al. Generation of a functional mammary gland from a single stem cell 2006;439.

[11] Van Keymeulen A, Rocha AS, Ousset M, Beck B, Bouvencourt G, Rock J, et al. Distinct stem cells contribute to mammary gland development and maintenance. Nature 2011;479:189–93.

[12] Morel AP, Ginestier C, Pommier RM, Cabaud O, Ruiz E, Wicinski J, et al. A stemness-related ZEB1-MSRB3 axis governs cellular pliancy and breast cancer genome stability. Nat Med 2017;23:568–78.

[13] Molyneux G, Geyer FC, Magnay FA, McCarthy A, Kendrick H, Natrajan R, et al. BRCA1 basal-like breast cancers originate from luminal epithelial progenitors and not from basal stem cells. Cell Stem Cell 2010;7:403–17.

[14] Li Y, Welm B, Podsypanina K, Huang S, Chamorro M, Zhang X, et al. Evidence that transgenes encoding components of the Wnt signaling pathway preferentially induce mammary cancers from progenitor cells. Proc Natl Acad Sci U S A 2003;100:15853–8.

[15] Lim E, Vaillant F, Wu D, Forrest NC, Pal B, Hart AH, et al. Aberrant luminal progenitors as the candidate target population for basal tumor development in BRCA1 mutation carriers. Nat Med 2009.

[16] Ginestier C, Wicha MS. Mammary stem cell number as a determinate of breast cancer risk. Breast Cancer Res 2007;9:109.

[17] Rios AC, Fu NY, Lindeman GJ, Visvader JE. In situ identification of bipotent stem cells in the mammary gland. Nature 2014.

[18] Rodilla V, Dasti A, Huyghe M, Lafkas D, Laurent C, Reyal F, et al. Luminal Progenitors Restrict Their Lineage Potential during Mammary Gland Development. PLoS Biol 2015;13.

[19] Janke R, Dodson AE, Rine J. Metabolism and Epigenetics. Annu Rev Cell Dev Biol 2015.

[20] Avgustinova A, Benitah SA. Epigenetic control of adult stem cell function. Nat Rev Mol Cell Biol 2016.

[21] Kanwal R, Gupta S. Epigenetic modifications in cancer. Clin Genet 2012.

[22] Stingl J, Eirew P, Ricketson I, Shackleton M, Vaillant F, Choi D, et al. Purification and unique properties of mammary epithelial stem cells. Nature 2006.

[23] Dontu G, Abdallah WM, Foley JM, Jackson KW, Clarke MF, Kawamura MJ, et al. In vitro propagation and transcriptional profiling of human mammary stem/progenitor cells. Genes Dev 2003;17:1253–70.

[24] Shaw FL, Harrison H, Spence K, Ablett MP, Simões BM, Farnie G, et al. A detailed mammosphere assay protocol for the quantification of breast stem cell activity. J Mammary Gland Biol Neoplasia 2012.

[25] Rota LM, Lazzarino DA, Ziegler AN, LeRoith D, Wood TL. Determining mammosphere-forming potential: Application of the limiting dilution analysis. J Mammary Gland Biol Neoplasia 2012.

[26] Hu Y, Smyth GK. ELDA: Extreme limiting dilution analysis for comparing depleted and enriched populations in stem cell and other assays. J Immunol Methods 2009.

[27] Key TJ, Appleby PN, Reeves GK, Roddam AW, Helzlsouer KJ, Alberg AJ, et al. Insulin-like growth factor 1 (IGF1), IGF binding protein 3 (IGFBP3), and breast cancer risk: Pooled individual data analysis of 17 prospective studies. Lancet Oncol 2010.

[28] De Ostrovich KK, Lambertz I, Colby JKL, Tian J, Rundhaug JE, Johnston D, et al. aracrine overexpression of insulin-like growth factor-1 enhances mammary tumorigenesis in vivo. Am J Pathol 2008.

[29] Bai L, Rohrschneider LR. s-SHIP promoter expression marks activated stem cells in developing mouse mammary tissue. Genes Dev 2010.

[30] Sitar T, Popowicz GM, Siwanowicz I, Huber R, Holak TA. Structural basis for the inhibition of insulin-like growth factors by insulin-like growth factor-binding proteins. Proc Natl Acad Sci U S A 2006.

[31] Allard JB, Duan C. IGF-binding proteins: Why do they exist and why are there so many? Front Endocrinol (Lausanne) 2018.

[32] Allar MA, Wood TL. Expression of the insulin-like growth factor binding proteins during postnatal development of the murine mammary gland. Endocrinology 2004.

[33] Richards RG, Klotz DM, Walker MP, Diaugustine RP. Mammary gland branching morphogenesis is diminished in mice with a deficiency of insulin-like growth factor-I (IGF-I), but not in mice with a liver-specific deletion of IGF-I. Endocrinology 2004;145:3106–10.

[34] Richert MM, Wood TL. The insulin-like growth factors (IGF) and IGF type I receptor during postnatal growth of the murine mammary gland: sites of messenger ribonucleic acid expression and potential functions. Endocrinology 1999.

[35] Lillycrop KA, Burdge GC. Maternal diet as a modifier of offspring epigenetics. J Dev Orig Health Dis 2015.

[36] Curradi M, Izzo A, Badaracco G, Landsberger N. Molecular Mechanisms of Gene Silencing Mediated by DNA Methylation. Mol Cell Biol 2002.

[37] Kikuchi K, Bichell DP, Rotwein P. Chromatin changes accompany the developmental activation of insulin-like growth factor I gene transcription. J Biol Chem 1992;267:21505–11.

[38] Oberbauer AM. The regulation of IGF-1 gene transcription and splicing during development and aging. Front Endocrinol (Lausanne) 2013.

[39] Lühr I, Friedl A, Overath T, Tholey A, Kunze T, Hilpert F, et al. Mammary fibroblasts regulate morphogenesis of normal and tumorigenic breast epithelial cells by mechanical and paracrine signals. Cancer Lett 2012.

[40] Bu W, Liu Z, Jiang W, Nagi C, Huang S, Edwards DP, et al. Mammary precancerous stem and non-stem cells evolve into cancers of distinct subtypes. Cancer Res 2019.

[41] Keller PJ, Arendt LM, Skibinski A, Logvinenko T, Klebba I, Dong S, et al. Defining the cellular precursors to human breast cancer. Proc Natl Acad Sci U S A 2012.

[42] Suvà ML, Riggi N, Bernstein BE. Epigenetic reprogramming in cancer. Science (80-) 2013.

[43] Ma M, Zhou QJ, Xiong Y, Li B, Li XT. Preeclampsia is associated with hypermethylation of IGF-1 promoter mediated by DNMT1. Am J Transl Res 2018.

[44] Ouni M, Gunes Y, Belot MP, Castell AL, Fradin D, Bougnères P. The igf 1 p2 promoter is an epigenetic qtl for circulating igf 1 and human growth. Clin Epigenetics 2015.

[45] Fung CM, Yang Y, Fu Q, Brown AS, Yu B, Callaway CW, et al. IUGR prevents IGF-1 upregulation in juvenile male mice by perturbing postnatal IGF-1 chromatin remodeling. Pediatr Res 2015;78:14–23.

[46] Ouni M, Gunes Y, Belot MP, Castell AL, Fradin D, Bougnères P. The igf 1 p2 promoter is an epigenetic qtl for circulating igf 1 and human growth. Clin Epigenetics 2015;7:2–12.

[47] Zhou J, Sears RL, Xing X, Zhang B, Li D, Rockweiler NB, et al. Tissue-specific DNA methylation is conserved across human, mouse, and rat, and driven by primary sequence conservation. BMC Genomics 2017.

[48] Rotwein P. Diversification of the insulin-like growth factor 1 gene in mammals. PLoS One 2017.

[49] Kim M, Costello J. DNA methylation: An epigenetic mark of cellular memory. Exp Mol Med 2017.

[50] Reis-Filho JS, Pusztai L. Gene expression profiling in breast cancer: Classification, prognostication, and prediction. Lancet 2011.

[51] Aupperlee MD, Zhao Y, Tan YS, Zhu Y, Langohr IM, Kirk EL, et al. Puberty-specific promotion of mammary tumorigenesis by a high animal fat diet. Breast Cancer Res 2015.

[52] Bu W, Liu Z, Jiang W, Nagi C, Huang S, Edwards DP, et al. Mammary precancerous stem and non-stem cells evolve into cancers of distinct subtypes. Cancer Res 2019.

[53] Van Hoeven KH, Drudis T, Cranor ML, Erlandson RA, Rosen PP. Low-grade adenosquamous carcinoma of the breast: A clinicopathologic study of 32 cases with ultrastructural analysis. Am J Surg Pathol 1993.

[54] Hollern DP, Swiatnicki MR, Andrechek ER. Histological subtypes of mouse mammary tumors reveal conserved relationships to human cancers. PLoS Genet 2018.

[55] Keller PJ, Arendt LM, Skibinski A, Logvinenko T, Klebba I, Dong S, et al. Defining the cellular precursors to human breast cancer. Proc Natl Acad Sci 2012.

[56] Slaga TJ. SENCAR mouse skin tumorigenesis model versus other strains and stocks of mice. Environ Health Perspect 1986.

[57] Kumaki Y, Oda M, Okano M. QUMA: quantification tool for methylation analysis. Nucleic Acids Res 2008.

[58] Cardiff RD, Anver MR, Gusterson BA, Hennighausen L, Jensen RA, Merino MJ, et al. The mammary pathology of genetically engineered mice: The consensus report and recommendations from the Annapolis meeting. Oncogene 2000.

