## Supplementary figures and images for "Early Dietary Exposures Epigenetically Program Mammary Cancer Susceptibility through IGF1-mediated Expansion of Mammary Stem Cells"

### Supplemental Figures

Zheng et al. Supplementary Fig. S1

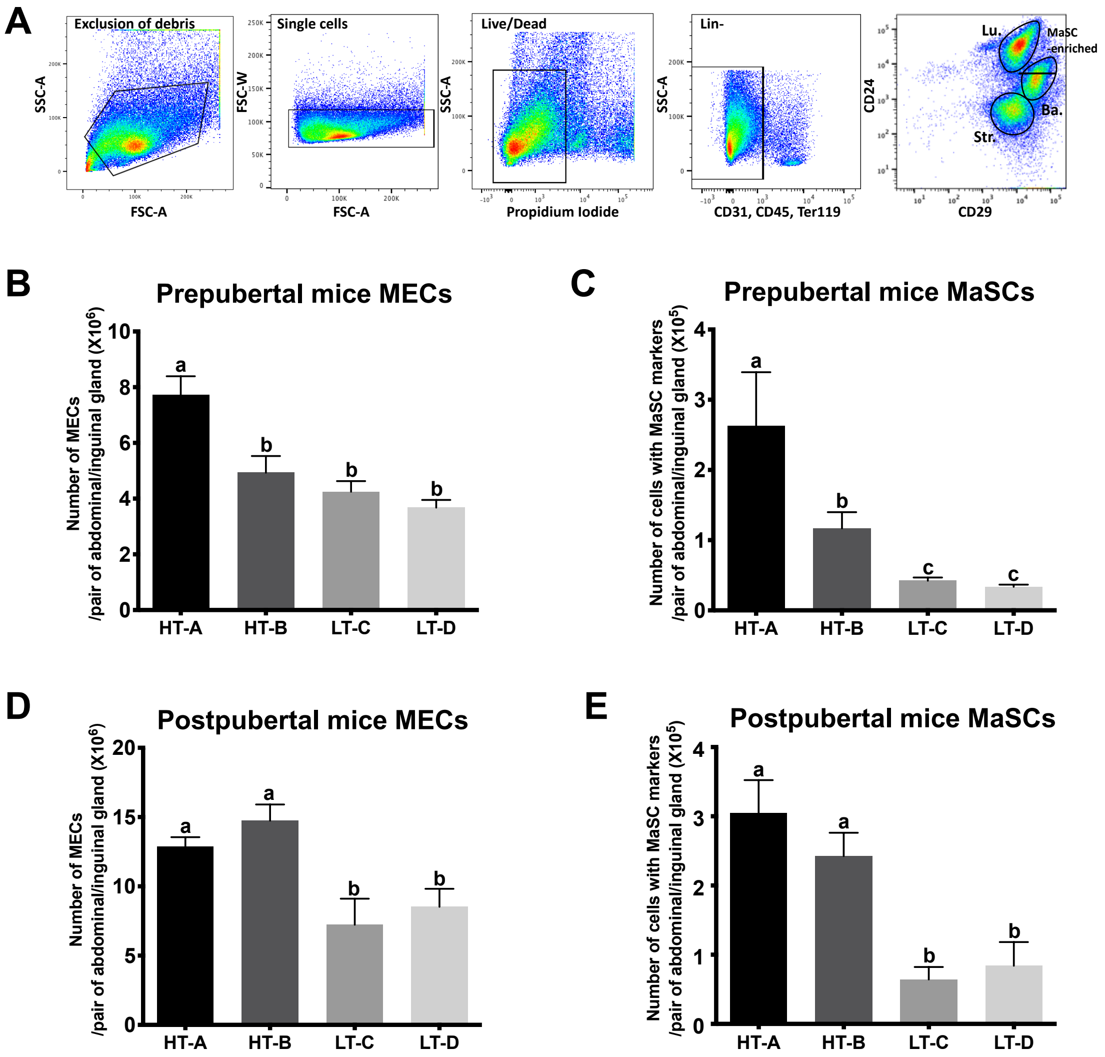

Zheng and Luo et al. Supplementary Fig. S2

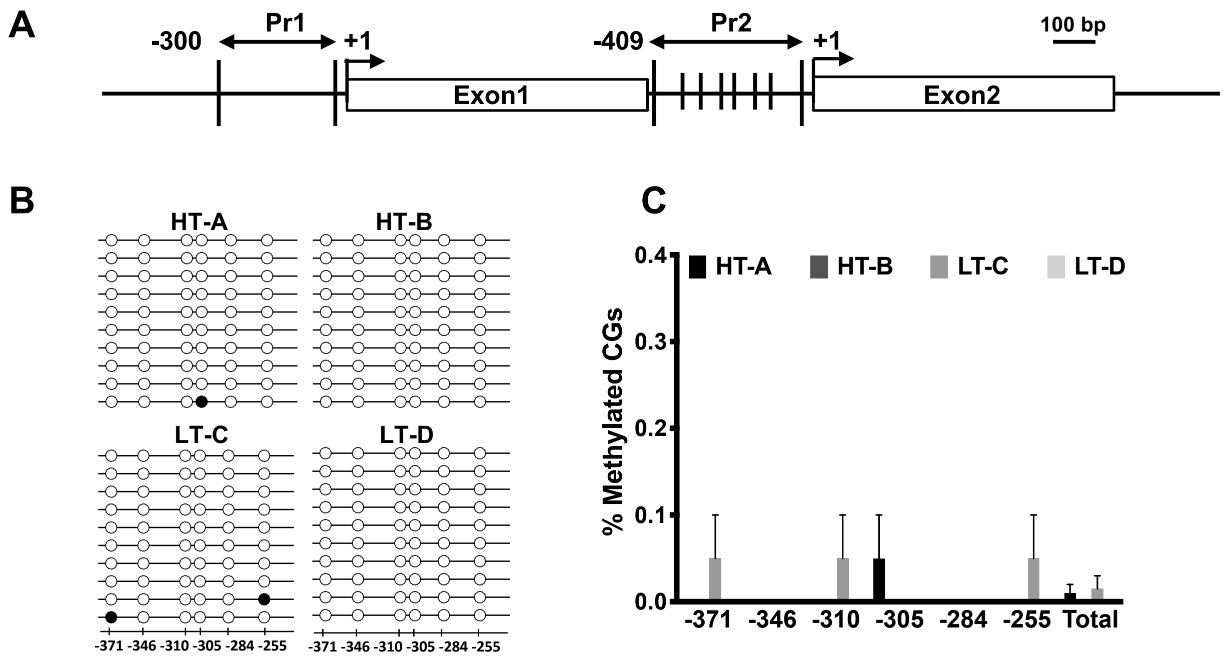
