## Supplemental Figure Legends for "Early Dietary Exposures Epigenetically Program Mammary Cancer Susceptibility through IGF1-mediated Expansion of Mammary Stem Cells"

### Supplementary Figure Legends

**Supplementary Figure S1. The absolute number of mammary epithelial cells with stem cell surface markers was increased in HT groups compared to LT groups. (A)** Gating strategy for sorting the MaSC-enriched population and the mammary stromal cell population from mammary glands**. (B)** Total numbers of rudimentary MECs harvested from prepubertal abdominal/inguinal mammary glands (n≥7) from HT and LT groups. **(C)** Estimates of the absolute number of cells with MaSC markers in prepubertal mice from HT and LT groups. The frequency of the MaSC-enriched population was multiplied by the number of MECs isolated from each corresponding sample. **(D)** Total numbers of rudimentary MECs harvested from postpubertal animals. **(E)** Estimates of the absolute number of cells with MaSC markers in postpubertal mice from HT and LT groups. Mean ± SEM are shown. One-way ANOVA was used for statistical analysis. Pairwise comparisons were performed using Fisher's LSD test. Columns are significantly different from each other if they do not share a letter (a≠b≠c, $P<0.05$).

**Supplementary Figure S2. IGF1 Pr2 was not differentially methylated in mammary stromal cells from HT and LT groups. (A)** Graphical representation of the IGF1 promoter 2 (Pr2) of the mouse with transcription start sites (TSSs) shown as arrows. CG sites used for the analysis shown as vertical bars. **(B)** DNA methylation patterns of representative animals from HT and LT groups. Each circle represents a CG site shown in panel (A), and each line represents a single clone, closed circles show methylated CG sites. **(C)** Quantification of DNA methylation percentages for each CG site (n=2).
