## Supplemental Materials and Methods for "Early Dietary Exposures Epigenetically Program Mammary Cancer Susceptibility through IGF1-mediated Expansion of Mammary Stem Cells"

### Supplementary Materials and Methods

#### Mammary Epithelial Cell Isolation

Fourth and fifth mammary gland pairs were resected from mice and minced with sterile scalpels until glands were rendered to a paste. The paste was dissociated using collagenase/hyaluronidase (Stem Cell Technologies, cat #07919) in a shaking incubator at 37°C, 230 rpm for 3 hours according to the manufacturer's protocol. After dissociation, cell pellets were collected by centrifugation. Red blood cells were lysed by washing the pellet in a 4:1 (v/v) mixture of 0.8% ammonium chloride ($NH_4Cl$) and cold Hanks' Balanced Salt Solution (HBSS, GE Healthcare Life Sciences, cat# SH30268.02) supplemented with 2% fetal bovine serum (FBS). A single-cell suspension was obtained by pipetting the cell pellet in 2ml of 0.25% Trypsin/EDTA solution (Stem Cell Technologies, cat #07901) for 1-2 minutes followed by the treatment with 5mg/mL of dispase (Gibco, cat#17105-041) supplemented with 10U of DNase I. The cell suspension was filtered through a 40μM mesh cell strainer.

#### Flow Cytometry and Cell Sorting

Cells from mammary gland dissociation were stained for 30 minutes at 4°C with fluorophore-conjugated antibodies at the specified concentrations shown in the Supplementary Table S4. After staining, cells were washed with DPBS three times and resuspended in 1mL of HBSS solution supplemented with 2% FBS. Propidium iodide (PI) (5ug/ml, Sigma, cat#P4147) was loaded into the cell suspension immediately before analysis, separating live and dead cells. CompBeads (BD BioSciences, cat#552845) were used as single color and unstained controls. For analysis of epithelial populations, stained cells were loaded onto a Fortessa X20 flow cytometer (BD

Biosciences). For each sample, 100,000 events were recorded, and data analysis was performed using the FlowJo software V10.4 (Tree Star.Inc.). For sorting of mammary stromal cells, a FACSAria II flow cytometer (BD Biosciences) was used. Sorted cell populations were reanalyzed and found to be 94%-98% pure, and cell viability was above 85%.

**Mammosphere Culture and Limiting Dilution Assay**

Cells freshly dissociated from the fourth and fifth pairs of mammary glands were plated in EpiCult™-B Basal Medium supplemented with 5% FBS, 10% EpiCult™-B Proliferation Supplement (Stem Cell Technologies, cat#05610), Human Epidermal Growth Factor (EGF, 10μg/mL, SC Tech, cat #78006), Recombinant Human Basic Fibroblast Growth Factor (FGF, 10μg/mL, SC Tech, cat #78003) and Heparin (50μg/mL, SC Tech, cat#07980) for 24 hours, followed by an additional 24 hours in serum-free EpiCult™-B Basal Medium. MECs were then trypsinized and plated at 10,000 cells/well in pHEMA-coated (Santa Cruz Biotechnology, cat# sc-253284) 24-well plates in DMEM/Ham's F-12 (Caisson Labs, cat#DFP17) supplemented with 1X B27 supplement (Gibco, cat#17504044), EGF (10μg/mL) and FGF (10μg/mL). After seven days, mammospheres with a diameter larger than 50μM were counted and collected. Mammospheres were dissociated to single cells by pipetting in a 0.25% Trypsin/EDTA solution (Stem Cell Technologies, cat#07901). Single cells from P1 mammospheres were then plated for P2 mammosphere formation at the same conditions described above. After seven days, P2 mammospheres were collected, counted and dissociated again into single cells as described above. Single cells from P2 mammospheres were plated into a 96-well ultra-low attachment plate at dilutions ranging from 1 to 512 cells per well (8 replicates per dilution). After seven days, wells were scored for the presence or absence of mammospheres. To assess the frequency of mammosphere-initiating cells, an extreme limiting dilution analysis was performed as described previously [23]. Pairwise differences between the groups were compared with likelihood ratio tests using the asymptotic chi-squared ($\chi 2$) test approximation to the log–ratio [23].

For performing conditioned media assays, conditioned media from mammary stromal cell cultures were diluted to a 50% concentration with DMEM/Ham's F-12. B27, EGF and FGF were added to the conditioned media with the same final concentrations described above. Single cells from P2 mammospheres were plated at 10,000 cells/well in pHEMA-coated 24-well plates. Recombinant IGF1 (Life Technologies, cat# PHG0071) and picropodophyllin (PPP, Santa Cruz, cat# sc-204008) were added to the conditioned media at a final concentration of 7.5 nM and 10 μM, respectively.

**RNA Extraction and RT-qPCR**

Total RNA was extracted from homogenized fourth and fifth pairs of mammary glands using the Quick-RNA™ Miniprep Plus Kit (Zymo Research, cat #R1057) following the manufacturer's protocol, and then reverse transcribed using M-MLV Reverse Transcriptase (Promega, cat #M1701) according to manufacturer's instructions. RT-qPCR was performed on an ABI Prism 7900HT sequence detection system (Applied Biosystems, Life Technologies Corporation, Carlsbad, CA). TaqMan assays were used to assess mammary gland mRNA levels of IGF1 (Assay ID: Mm00439560_m1, ThermoFisher Scientific), IGFBP5 (Assay ID: Mm00516037_m1) and TATA-binding protein (TBP) (Assay ID: Mm01277042_m1). All the other RT-qPCR reactions were performed with the PowerUp SYBR Green Master Mix (Applied Biosystem, cat #4367659) with primers listed in the Supplementary Table S5. Target gene expression was normalized to TBP.

All assays, including target and reference genes, were run as triplicates on the same plate. The $2^{-\Delta\Delta Ct}$ method was used to calculate relative gene expression levels.

**Western blot analysis**

Snap-frozen abdominal/inguinal mammary glands were extracted with boiling 2x Laemmli sample buffer. Protein concentrations were determined using the Pierce™ BCA protein assay kit (Thermo Fisher Scientific, Cat #23225). Protein samples (60μg) were resolved on SDS-PAGE gels and transferred onto 0.2μM PVDF membranes (EMD Millipore, cat #ISEQ00010). After blocking with 3% milk/TBST, membranes were probed with the primary antibodies, followed by the appropriate HRP-conjugated secondary antibody shown in Supplementary Table S4. Bands were detected using the ECL Plus kit (Amersham, GE Healthcare, Piscataway, NJ). Images were captured with a FluorChem M imager and quantified with the AlphaView SA software (ProteinSimple, San Jose, CA). Signals of target protein bands were normalized to GAPDH bands of the same sample and then normalized to the control group to calculate fold changes.
